# Genomic analyses for age at menarche identify 389 independent signals and indicate BMI-independent effects of puberty timing on cancer susceptibility

**DOI:** 10.1101/076794

**Authors:** Felix R. Day, Deborah J. Thompson, Hannes Helgason, Daniel I. Chasman, Hilary Finucane, Patrick Sulem, Katherine S. Ruth, Sean Whalen, Abhishek K. Sarkar, Eva Albrecht, Elisabeth Altmaier, Marzyeh Amini, Caterina M. Barbieri, Thibaud Boutin, Archie Campbell, Ellen Demerath, Ayush Giri, Chunyan He, Jouke J. Hottenga, Robert Karlsson, Ivana Kolcic, Po-Ru Loh, Kathryn L. Lunetta, Massimo Mangino, Brumat Marco, George McMahon, Sarah E. Medland, Ilja M. Nolte, Raymond Noordam, Teresa Nutile, Lavinia Paternoster, Natalia Perjakova, Eleonora Porcu, Lynda M. Rose, Katharina E. Schraut, Ayellet V. Segrè, Albert V. Smith, Lisette Stolk, Alexander Teumer, Irene L. Andrulis, Stefania Bandinelli, Matthias W. Beckmann, Javier Benitez, Sven Bergmann, Murielle Bochud, Eric Boerwinkle, Stig E. Bojesen, Manjeet K. Bolla, Judith S. Brand, Hiltrud Brauch, Hermann Brenner, Linda Broer, Thomas Brüning, Julie E. Buring, Harry Campbell, Eulalia Catamo, Stephen Chanock, Georgia Chenevix-Trench, Tanguy Corre, Fergus J. Couch, Diana L. Cousminer, Angela Cox, Laura Crisponi, Kamila Czene, George Davey-Smith, Eco J.C.N de Geus, Renée de Mutsert, Immaculata De Vivo, Joe Dennis, Peter Devilee, Isabel dos-Santos-Silva, Alison M. Dunning, Johan G. Eriksson, Peter A. Fasching, Lindsay Fernández-Rhodes, Luigi Ferrucci, Dieter Flesch-Janys, Lude Franke, Marike Gabrielson, Ilaria Gandin, Graham G. Giles, Harald Grallert, Daniel F. Gudbjartsson, Pascal Guénel, Per Hall, Emily Hallberg, Ute Hamann, Tamara B. Harris, Catharina A. Hartman, Gerardo Heiss, Maartje J. Hooning, John L. Hopper, Frank Hu, David Hunter, M. Arfan Ikram, Hae Kyung Im, Marjo-Riitta Järvelin, Peter K. Joshi, David Karasik, Zoltan Kutalik, Genevieve LaChance, Diether Lambrechts, Claudia Langenberg, Lenore J. Launer, Joop S.E. Laven, Stefania Lenarduzzi, Jingmei Li, Penelope A. Lind, Sara Lindstrom, YongMei Liu, Jian'an Luan, Reedik Mägi, Arto Mannermaa, Hamdi Mbarek, Mark I. McCarthy, Christa Meisinger, Thomas Meitinger, Cristina Menni, Andres Metspalu, Kyriaki Michailidou, Lili Milani, Roger L. Milne, Grant W. Montgomery, Anna M. Mulligan, Mike A. Nalls, Pau Navarro, Heli Nevanlinna, Dale R. Nyholt, Albertine J. Oldehinkel, Tracy A. O'Mara, Aarno Palotie, Nancy Pedersen, Annette Peters, Julian Peto, Paul D.P. Pharoah, Anneli Pouta, Paolo Radice, Iffat Rahman, Susan M. Ring, Antonietta Robino, Frits R. Rosendaal, Igor Rudan, Rico Rueedi, Daniela Ruggiero, Cinzia F. Sala, Marjanka K. Schmidt, Robert A. Scott, Mitul Shah, Rossella Sorice, Melissa C. Southey, Ulla Sovio, Meir Stampfer, Maristella Steri, Konstantin Strauch, Toshiko Tanaka, Emmi Tikkanen, Nicholas J. Timpson, Michela Traglia, Thérèse Truong, Jonathan P. Tyrer, André G. Uitterlinden, Digna R. Velez Edwards, Veronique Vitart, Uwe Völker, Peter Vollenweider, Qin Wang, Elisabeth Widen, Ko Willems van Dijk, Gonneke Willemsen, Robert Winqvist, Bruce H.R Wolffenbuttel, Jing Hua Zhao, Magdalena Zoledziewska, Marek Zygmunt, Behrooz Z. Alizadeh, Dorret I. Boomsma, Marina Ciullo, Francesco Cucca, Tõnu Esko, Nora Franceschini, Christian Gieger, Vilmundur Gudnason, Caroline Hayward, Peter Kraft, Debbie A. Lawlor, Patrik K.E Magnusson, Nicholas G. Martin, Dennis O. Mook-Kanamori, Ellen A. Nohr, Ozren Polasek, David Porteous, Alkes L. Price, Paul M. Ridker, Harold Snieder, Tim D. Spector, Doris Stöckl, Daniela Toniolo, Sheila Ulivi, Jenny A. Visser, Henry Völzke, Nicholas J. Wareham, James F. Wilson, The LifeLines Cohort Study, The InterAct Consortium, kConFab/AOCS Investigators, Endometrial Cancer Association Consortium, Ovarian Cancer Association Consortium, PRACTICAL consortium, Amanda B. Spurdle, Unnur Thorsteindottir, Katherine S. Pollard, Douglas F. Easton, Joyce Y. Tung, Jenny Chang-Claude, David Hinds, Anna Murray, Joanne M. Murabito, Kari Stefansson, Ken K. Ong, John R.B Perry

**Affiliations:** MRC Epidemiology Unit, University of Cambridge School of Clinical Medicine, Box 285 Institute of Metabolic Science, Cambridge Biomedical Campus, Cambridge, CB2 0QQ, UK; Centre for Cancer Genetic Epidemiology, Department of Public Health and Primary Care, University of Cambridge, CB1 8RN, UK; deCODE genetics/Amgen, Inc., IS-101 Reykjavik, Iceland; School of Engineering and Natural Sciences, University of Iceland, IS-101 Reykjavik, Iceland; Division of Preventive Medicine, Brigham and Women’s Hospital, Boston, MA 02215; Harvard Medical School, Boston, MA 02115, USA; Department of Epidemiology, Harvard School of Public Health, Boston, MA 02115, USA; Department of Mathematics, Massachusetts Institute of Technology, Cambridge, Massachusetts 02139-4307, USA; Genetics of Complex Traits, University of Exeter Medical School, University of Exeter, Exeter, EX2 5DW, UK; Gladstone Institutes, San Francisco, California, 94158, USA; Computer Science and Artificial Intelligence Lab, Massachusetts Institute of Technology, Cambridge, MA, USA; Broad Institute of the Massachusetts Institute of Technology and Harvard University, 140 Cambridge 02142, MA, USA; Institute of Genetic Epidemiology, Helmholtz Zentrum München – German Research Center for Environmental Health, 85764 Neuherberg, Germany; Institute of Epidemiology II, Helmholtz Zentrum München – German Research Center for Environmental Health, 85764 Neuherberg, Germany; Research Unit of Molecular Epidemiology, Helmholtz Zentrum München – German Research Center for Environmental Health, 85764 Neuherberg, Germany; Department of Epidemiology, University of Groningen, University Medical Center Groningen, Groningen, The Netherlands; Genetics of Common Disorders Unit, IRCCS San Raffaele Scientific Institute and Vita-Salute San Raffaele University, Milan, Italy; Medical Research Council Human Genetics Unit, Institute of Genetics and Molecular Medicine, University of Edinburgh, Edinburgh EH4 2XU, UK; Medical Genetics Section, Centre for Genomic and Experimental Medicine, Institute of Genetics and Molecular Medicine, University of Edinburgh, Edinburgh EH4 2XU, UK; Division of Epidemiology & Community Health, University of Minnesotta, Minneapolis MN 55455; Division of Epidemiology, Institute for Medicine and Public Health, Vanderbilt University, Nashville, TN 37235, USA; Vanderbilt Genetics Institute, Vanderbilt University, Nashville, TN; Department of Epidemiology, Indiana University Richard M. Fairbanks School of Public Health, Indianapolis, IN 46202, USA; Indiana University Melvin and Bren Simon Cancer Center, Indianapolis, IN 46202, USA; Department of Biological Psychology, VU University Amsterdam, van der Boechorststraat 1, 1081 BT, Amsterdam, The Netherlands; Department of Medical Epidemiology and Biostatistics, Karolinska Institutet, 17177 Stockholm, Sweden; Faculty of Medicine, University of Split, Split, Croatia; Program in Medical and Population Genetics, Broad Institute, Cambridge, MA, USA; Boston University School of Public Health, Department of Biostatistics. Boston, Massachusetts 02118, USA; NHLBI's and Boston University's Framingham Heart Study, Framingham, Massachusetts 01702-5827, USA; Department of Twin Research and Genetic Epidemiology, King's College London, London SE1 7EH, UK; National Institute for Health Research (NIHR) Biomedical Research Centre at Guy’s and St. Thomas’ Foundation Trust, London, UK; Department of Clinical Medical Sciences, Surgical and Health, University of Trieste, 34149 Trieste, Italy; School of Social and Community Medicine, University of Bristol, Bristol BS8 2BN, UK; QIMR Berghofer Medical Research Institute, Brisbane, Queensland, Australia; Department of Internal Medicine, Section Gerontology and Geriatrics, Leiden University Medical Center, Leiden, the Netherlands; Institute of Genetics and Biophysics – CNR, via Pietro Castellino 111, 80131, Naples, Italy; MRC Integrative Epidemiology Unit, University of Bristol, Bristol, UK; Estonian Genome Center, University of Tartu, Tartu, 51010, Estonia; Institute of Genetics and Biomedical Research, National Research Council, Cagliari, 09042 Sardinia, Italy; Centre for Cardiovascular Sciences, Queen's Medical Research Institute, University of Edinburgh, Royal Infirmary of Edinburgh, Little France Crescent, Edinburgh, EH16 4TJ, Scotland; Centre for Global Health Research, Usher Institute of Population Health Sciences and Informatics, University of Edinburgh, Teviot Place, Edinburgh, EH8 9AG, Scotland; Cancer Program, Broad Institute, Cambridge, MA, USA; Faculty of Medicine, University of Iceland, IS-101 Reykjavik, Iceland; Icelandic Heart Association, Kopavogur, Iceland; Department of Internal Medicine, Erasmus MC, 3015GE Rotterdam, the Netherlands; Institute for Community Medicine, University Medicine Greifswald, 17475 Greifswald, Germany; Fred A. Litwin Center for Cancer Genetics, Lunenfeld-Tanenbaum Research Institute of Mount Sinai Hospital, Toronto, ON, Canada; Department of Molecular Genetics, University of Toronto, Toronto, Ontario, Canada; Geriatric Unit, Azienda Sanitaria di Firenze, Florence, Italy; Department of Gynaecology and Obstetrics, University Hospital Erlangen, Friedrich-Alexander University Erlangen-Nuremberg, Erlangen, Germany; Human Genetics Group, Human Cancer Genetics Program, Spanish National Cancer Research Centre (CNIO), Madrid, Spain; Centro de Investigación en Red de Enfermedades Raras (CIBERER), Valencia, Spain; Swiss Institute of Bioinformatics, CH-1015, Lausanne, Switzerland; Department of Computational Biology, University of Lausanne, Lausanne, Switzerland; Institute of Social and Preventive Medicine, University Hospital of Lausanne, Lausanne, Switzerland; Human Genetics Center, School of Public Health, The University of Texas Health Science Center at Houston, Houston, TX 77030, USA; Copenhagen General Population Study, Herlev Hospital, Copenhagen University Hospital, University of Copenhagen, Copenhagen, Denmark; Department of Clinical Biochemistry, Herlev Hospital, Copenhagen University Hospital, University of Copenhagen, Copenhagen, Denmark; Faculty of Health and Medical Sciences, University of Copenhagen, Copenhagen, Denmark; Dr. Margarete Fischer-Bosch-Institute of Clinical Pharmacology, Stuttgart, Germany; University of Tübingen, Tübingen, Germany; German Cancer Consortium (DKTK), German Cancer Research Center (DKFZ), Heidelberg, Germany; Division of Clinical Epidemiology and Aging Research, German Cancer Research Center (DKFZ), Heidelberg, Germany; Division of Preventive Oncology, German Cancer Research Center (DKFZ) and National Center for Tumor Diseases (NCT), Heidelberg, Germany; Institute for Prevention and Occupational Medicine of the German Social Accident Insurance, Institute of the Ruhr University Bochum (IPA), Bochum, Germany; Institute for Maternal and Child Health – IRCCS “Burlo Garofolo”, 34137 Trieste, Italy; Division of Cancer Epidemiology and Genetics, National Cancer Institute, Bethesda, MD, USA; Department of Genetics, QIMR Berghofer Medical Research Institute, Brisbane, Australia; Department of Laboratory Medicine and Pathology, Mayo Clinic, Rochester, MN, USA; Division of Genetics, Children’s Hospital of Philadelphia, Philadelphia, PA, USA; Department of Genetics, University of Pennsylvania, Philadelphia, PA, USA; Academic Unit of Molecular Oncology, Department of Oncology and Metabolism, University of Sheffield, Sheffield, UK; Department of Clinical Epidemiology, Leiden University Medical Center, Leiden, the Netherlands; Channing Division of Network Medicine, Department of Medicine, Brigham and Women’s Hospital and Harvard Medical School, Boston, MA 02115, USA; Department of Pathology, Leiden University Medical Center, Leiden, The Netherlands; Department of Human Genetics, Leiden University Medical Center, 2300 RC Leiden, The Netherlands; Non-communicable Disease Epidemiology Department, London School of Hygiene and Tropical Medicine, London, UK; Centre for Cancer Genetic Epidemiology, Department of Oncology, University of Cambridge, Cambridge, CB1 8RN, UK; Department of General Practice and Primary health Care, University of Helsinki, Finland; David Geffen School of Medicine, Department of Medicine Division of Hematology and Oncology, University of California at Los Angeles, CA, USA; Department of Epidemiology, Gillings School of Global Public Health, University of North Carolina, Chapel Hill, NC 27514; Longitudinal Studies Section, Translational Gerontology Branch, National Institute on Aging, Baltimore, Maryland 21224, United States of America; Institute for Medical Biometrics and Epidemiology, University Clinic Hamburg-Eppendorf, Hamburg, Germany; Department of Cancer Epidemiology/Clinical Cancer Registry, University Clinic Hamburg-Eppendorf, Hamburg, Germany; Department of Genetics, University of Groningen, University Medical Centre Groningen, Groningen, The Netherlands; Cancer Epidemiology Centre, Cancer Council Victoria, Melbourne, Australia; Centre for Epidemiology and Biostatistics, Melbourne School of Population and Global Health, The University of Melbourne, Melbourne, Australia; German Center for Diabetes Research, 85764 Neuherberg, Germany; Cancer & Environment Group, Center for Research in Epidemiology and Population Health (CESP), INSERM, University Paris-Sud, University Paris-Saclay, Villejuif, France; Division of Epidemiology, Department of Health Sciences Research, Mayo Clinic, Rochester, Minnesota, USA; Molecular Genetics of Breast Cancer, Deutsches Krebsforschungszentrum (DKFZ), Heidelberg, Germany; Laboratory of Epidemiology and Population Sciences, National Institute on Aging, Intramural Research Program, National Institutes of Health, Bethesda, Maryland, 20892, USA; Department of Psychiatry, University of Groningen, University Medical Center Groningen, Groningen, The Netherlands; Department of Medical Oncology, Family Cancer Clinic, Erasmus MC Cancer Institute, Rotterdam, The Netherlands; Department of Nutrition, Harvard School of Public Health, Boston, MA 02115, USA; Department of Epidemiology, Erasmus MC, Rotterdan, the Netherlands; Section of Genetic Medicine, Department of Medicine, University of Chicago, Chicago, IL, USA; Department of Epidemiology and Biostatistics, MRC Health Protection Agency (HPA) Centre for Environment and Health, School of Public Health, Imperial College London, UK; Biocenter Oulu, P.O.Box 5000, Aapistie 5A, FI-90014 University of Oulu, Finland; Department of Children and Young People and Families, National Institute for Health and Welfare, Aapistie 1, Box 310, FI-90101 Oulu, Finland; Institute of Health Sciences, P.O.Box 5000, FI-90014 University of Oulu, Finland; Unit of Primary Care, Oulu University Hospital, Kajaanintie 50, P.O.Box 20, FI-90220 Oulu, 90029 OYS, Finland; Hebrew SeniorLife Institute for Aging Research, Boston, MA, 02131, USA; Laboratory for Translational Genetics, Department of Oncology, University of Leuven, Leuven, Belgium; Vesalius Research Center (VRC), VIB, Leuven, Belgium; Division of Reproductive Medicine, Department of Obstetrics and Gynaecology, Erasmus MC, Rotterdam, The Netherlands; Department of Epidemiology, School of Public Health, University of Washington, Seattle, WA 98195, USA; Center for Human Genetics, Division of Public Health Sciences, Wake Forest School of Medicine; Translational Cancer Research Area, University of Eastern Finland, Kuopio, Finland; Institute of Clinical Medicine, Pathology and Forensic Medicine, University of Eastern Finland, Kuopio, Finland; Imaging Center, Department of Clinical Pathology, Kuopio University Hospital, Kuopio, Finland; NIHR Oxford Biomedical Research Centre, Churchill Hospital, OX3 7LE Oxford, UK; Oxford Centre for Diabetes, Endocrinology, & Metabolism, University of Oxford, Churchill Hospital, OX3 7LJ Oxford, UK; Wellcome Trust Centre for Human Genetics, University of Oxford, Oxford, UK; Central Hospital of Augsburg, MONICA/KORA Myocardial Infarction Registry, Augsburg, Germany; Institute of Human Genetics, Helmholtz Zentrum München, German Research Center for Environmental Health, Neuherberg, Germany; Department of Electron Microscopy/Molecular Pathology, The Cyprus Institute of Neurology and Genetics, Nicosia, Cyprus; Institute for Molecular Bioscience, The University of Queensland, Brisbane, Australia; Department of Laboratory Medicine and Pathobiology, University of Toronto, Toronto, ON, Canada; Laboratory Medicine Program, University Health Network, Toronto, ON, Canada; Laboratory of Neurogenetics, National Institute on Aging, Bethesda, MD, USA; Department of Obstetrics and Gynecology, Helsinki University Hospital, University of Helsinki, Helsinki, Finland; Institute of Health and Biomedical Innovation, Queensland University of Technology, Australia; Interdisciplinary Center Psychopathology and Emotion Regulation, University of Groningen, University Medical Center Groningen, Groningen, The Netherlands; British Heart Foundation Glasgow Cardiovascular Research Centre, Institute of Cardiovascular and Medical Sciences, College of Medical, Veterinary and Life Sciences, University of Glasgow, Glasgow G12 8TA, UK; Psychiatric & Neurodevelopmental Genetics Unit, Department of Psychiatry, Massachusetts General Hospital, Boston, MA, USA; Stanley Center for Psychiatric Research, Broad Institute of MIT and Harvard, Cambridge, Massachusetts 02142, USA; Wellcome Trust Sanger Institute, Wellcome Trust Genome Campus, Hinxton, UK; Analytic and Translational Genetics Unit, Massachusetts General Hospital and Harvard Medical School, Boston, Massachusetts, USA; Institute for Molecular Medicine Finland (FIMM), University of Helsinki, Finland; National Institute for Health and Welfare, Finland; Unit of Molecular Bases of Genetic Risk and Genetic Testing, Department of Preventive and Predictive Medicine, Fondazione IRCCS Istituto Nazionale dei Tumori (INT), Milan, Italy; Institute of Environmental Medicine, Karolinska Institutet, Stockholm, Sweden; Division of Molecular Pathology, The Netherlands Cancer Institute – Antoni van Leeuwenhoek Hospital, Amsterdam, The Netherlands; Division of Psychosocial Research and Epidemiology, The Netherlands Cancer Institute – Antoni van Leeuwenhoek hospital, Amsterdam, The Netherlands; Department of Pathology, The University of Melbourne, Melbourne, Australia; Department of Obstetrics and Gynaecology, University of Cambridge, Cambridge, United Kingdom; Institute of Medical Informatics, Biometry and Epidemiology, Chair of Genetic Epidemiology, Ludwig-Maximilians-Universität, 81377 Munich, Germany; Department of Public Health, University of Helsinki, Helsinki, Finland; Vanderbilt Epidemiology Center, Institute for Medicine and Public Health, Vanderbilt University, Nashville, TN, USA; Department of Obstetrics and Gynecology, Vanderbilt University School of Medicine, Nashville, TN, USA; Interfaculty Institute for Genetics and Functional Genomics, University Medicine Greifswald, 17475 Greifswald, Germany; University Hospital of Lausanne, Lausanne, Switzerland; Department of Internal Medicine, Division of Endocrinology, Leiden University Medical Center, Leiden, the Netherlands; Einthoven Laboratory for Experimental Vascular Medicine, Leiden University Medical Center, Leiden, the Netherlands; Laboratory of Cancer Genetics and Tumor Biology, Cancer and Translational Medicine Research Unit, Biocenter Oulu, University of Oulu, Oulu, Finland; Laboratory of Cancer Genetics and Tumor Biology, Northern Finland Laboratory Centre NordLab, Oulu, Finland; Department of Endocrinology, University of Groningen, University Medical Centre Groningen, Groningen, The Netherlands; Department of Obstetrics and Gynecology, University Medicine Greifswald, 17475 Greifswald, Germany; University of Sassari, Department of Biomedical Sciences, Sassari, 07100 Sassari, Italy; Department of Biostatistics, Harvard School of Public Health, Boston, MA 02115, USA; Department of Public Health and Primary Care, Leiden University Medical Center, Leiden, the Netherlands; Research Unit for Gynaecology and Obstetrics, Department of Clinical Research, University of Southern Denmark, Denmark; Department of Obstetrics and Gynaecology, Campus Grosshadern, Ludwig-Maximilians-University, Munich, Germany; Full consortium membership is displayed in the supplementary material; Division of Biostatistics, Institute for Human Genetics, and Institute for Computational Health Sciences, University of California, San Francisco, California, 94158, USA; 23andMe Inc., 899 W. Evelyn Avenue, Mountain View, California 94041, USA; Division of Cancer Epidemiology, German Cancer Research Center (DKFZ), Heidelberg, Germany; University Cancer Center Hamburg (UCCH), University Medical Center Hamburg-Eppendorf, Hamburg, Germany; Boston University School of Medicine, Department of Medicine, Section of General Internal Medicine, Boston, MA 02118, USA; Department of Paediatrics, University of Cambridge,Cambridge, CB2 0QQ, UK

**Author notes:** denotes equal contribution (Felix R. Day, Deborah J. Thompson, Hannes Helgason, Kari Stefansson, Ken K. Ong, John R.B Perry). Correspondence to John R.B. Perry and Ken K. Ong.

## Abstract

The timing of puberty is a highly polygenic childhood trait that is epidemiologically associated with various adult diseases. Here, we analyse 1000-Genome reference panel imputed genotype data on up to ~370,000 women and identify 389 independent signals (all P<5×10^−8^) for age at menarche, a notable milestone in female pubertal development. In Icelandic data from deCODE, these signals explain ~7.4% of the population variance in age at menarche, corresponding to one quarter of the estimated heritability. We implicate over 250 genes via coding variation or associated gene expression, and demonstrate enrichment across genes active in neural tissues. We identify multiple rare variants near the imprinted genes *MKRN3* and *DLK1* that exhibit large effects on menarche only when paternally inherited. Disproportionate effects of variants on early or late puberty timing are observed: single variant and heritability estimates are larger for early than late puberty timing in females. The opposite pattern is seen in males, with larger estimates for late than early puberty timing. Mendelian randomization analyses indicate causal inverse associations, independent of BMI, between puberty timing and risks for breast and endometrial cancers in women, and prostate cancer in men. In aggregate, our findings reveal new complexity in the genetic regulation of puberty timing and support new causal links with adult cancer risks.

## Introduction

Puberty is the developmental stage of transition from childhood to physical and sexual maturity and its timing varies markedly between individuals^1^. This variation reflects the influence of genetic, nutritional and other environmental factors and is associated with the subsequent risks for several diseases in adult life^2^. Our previous large-scale genomic studies identified 113 independent regions associated with age at menarche (AAM), a well-recalled milestone of puberty in females^3,4^. The vast majority of those signals have concordant effects on the age at voice breaking, a corresponding milestone in males^5^. Those genetic findings implicated a diverse range of mechanisms involved in the regulation of puberty timing, identified significant enrichment of AAM-associated variants in/near genes disrupted in rare disorders of puberty, and highlighted shared aetiological factors between puberty timing and metabolic disease outcomes^2,3^.

However, those previous studies were based on genome-wide association data that were imputed to the relatively sparse HapMap2 reference panel or they used gene-centric arrays. Consequently, the reported genetic signals explained only a small fraction of the population variance, suggesting that several hundreds or thousands of signals are involved^3,4^. Here, we report an enlarged genomic analysis for AAM in a nearly 2-fold higher sample of women than previously^3^, and using more densely imputed genomic data. Our findings increase by more than 3-fold the number of independently associated signals and indicate likely causal effects of puberty timing on risks of various sex steroid sensitive cancers in men and women.

## Results

Genome-wide array data, imputed to the 1000-Genome reference panel, were available in up to 329,345 women of European ancestry. These comprised 40 studies from the ReproGen consortium (N=179,117), in addition to the 23andMe, Inc. (N=76,831) and UK Biobank studies (N=73,397) (**Table S1**), which combined demonstrated no evidence of test statistic inflation due to population structure (LD score intercept =1, s.e 0.02). In total, 37,925 variants were associated with AAM at P<5×10^−8^, which were resolved to 389 statistically-independent signals (Figure 1, **Table S2**). Per-allele effect sizes ranged from ~1 week to 5 months, 16 index variants were classed as low-frequency (minor allele frequency <5%; minimum observed 0.5%), and 26 were insertion/deletion polymorphisms. Signals were distributed evenly across all 23 chromosomes with respect to chromosome size (**Figure S1**). Of the previously reported 106 autosomal, 5 exome-array and 2 X-chromosome signals for AAM, all remained associated at genome-wide significance, except for two common loci (reported as *SCRIB*/*PARP10* [P=5x10^−4^] and *FUT8* [P=5.4x10^−7^]) and one rare variant not captured by the 1000G reference panel (p.W275X, *TACR3*).

**Figure 1.**
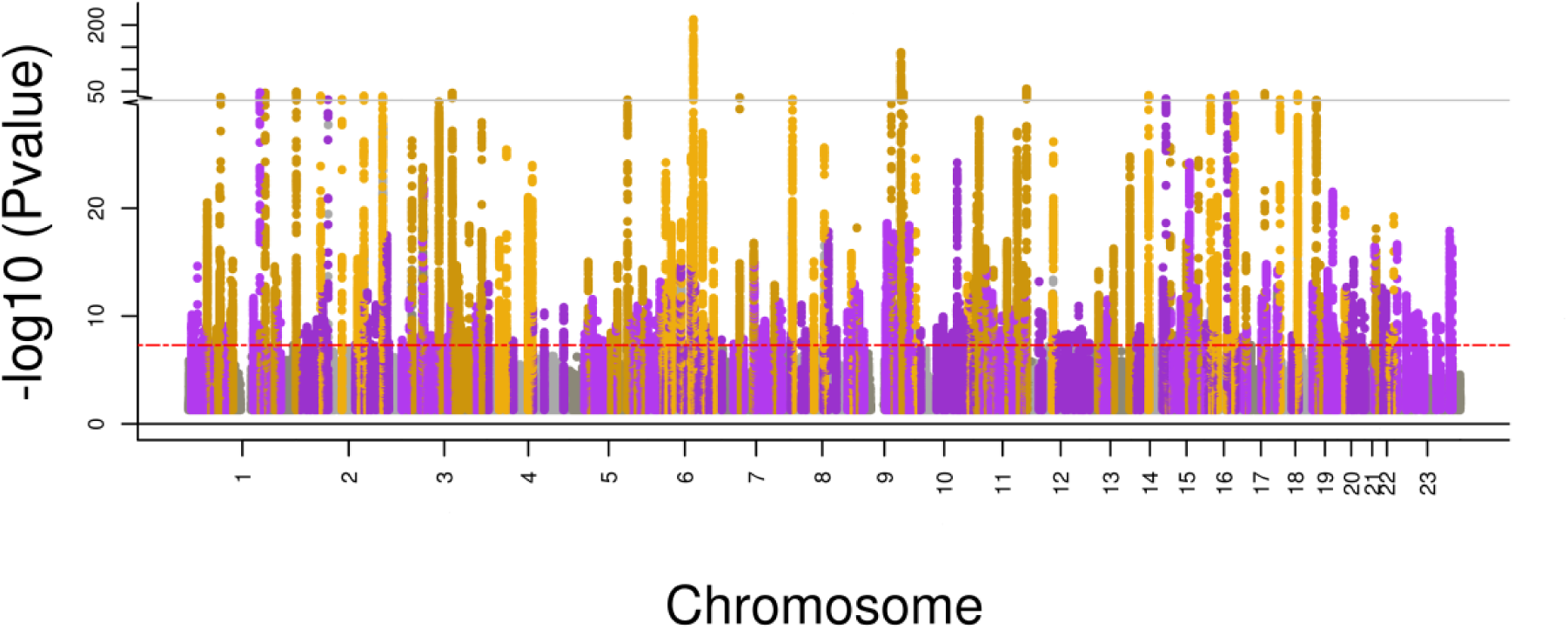
‘Manhattan’ plot for GWAS of age at menarche. Previously identified genome-wide significant loci are indicated in gold, novel loci are indicated in purple. SNPs within 300kb of the lead SNP at each loci are highlighted. The y-axis has been truncated above 30.

Independent replication in the deCODE study (N=39,543 women) showed that 367 (94.3%) of the 389 signals had directionally-concordant trends or associations with AAM (binomial P<4.0×10^−82^), and 187 signals reached nominal significance for association (P<0.05) (**Table S3**). In aggregate, the top 389 index SNPs explained 7.4% of the trait variance in deCODE and 7.2% in UK Biobank (using weights derived from a meta-analysis excluding UK Biobank). These estimates are double that of the previously reported 106 signals^3^ (3.7% in deCODE) and equivalent to one quarter of the total chip-captured heritability (h^2^ _SNP_=32%, se=1%) for AAM, estimated in UK Biobank.

Consistent with our previous reports, we found a strongly shared genetic architecture between AAM in women and age at voice breaking in men (considered as a continuous trait in 55,871 men in 23andMe, Inc.) (genetic correlation (rg)=0.75 P=1.2×10^−79^). Of the 389 AAM signals, 327 demonstrated directionally-consistent trends or associations with age at voice breaking in men (binomial P=1.4×10^−44^), and 127 signals reached nominal significance for association (i.e. P<0.05) (**Table S4**). Similarly, in UK Biobank where age at voice breaking was recorded using only 3 categories, 277 and 297 of the 377 autosomal loci demonstrated directionally-consistent trends or associations with “relatively early voice breaking” (N=2,678 cases, N=55,763 controls, binomial P=2.4×10^−20^) and “relatively late voice breaking” (N=3,566 cases, P=1.9×10^−30^), respectively (**Table S5**).

## Implicated genes and tissues

We used a number of analytical techniques to implicate genes in the regulation of AAM. These included: mapping of non-synonymous SNPs, gene expression QTLs and integration of Hi-C chromatin interaction data. Eight of the 389 lead variants were non-synonymous, and a further 24 genes were implicated by highly correlated non-synonymous variants (r^2^>0.8) (**Table S6**). These include genes disrupted in rare disorders of puberty: aromatase (*CYP19A1*, #307), gonadotropin-releasing hormone (*GNRH1*, #178), kisspeptin (*KISS1,* signal #31); and the stop-gained variant in fucosyltransferase 2 (*FUT2*, #357) that confers blood group secretor status.

Two approaches were used to interrogate publicly available gene expression datasets, both of which use one or more SNPs (not restricted to lead SNPs) to infer patterns of gene expression based on imputation reference panels (see **methods**). Firstly, under the hypothesis that some causal genes and regulatory mechanisms might be ubiquitously expressed, we analysed data from the largest available eQTL dataset for any tissue (whole blood, N=5,311)^6^. Systematic eQTL integration using the Summary Mendelian Randomization approach^7^ prioritised 113 transcripts, for 60 of which there was evidence for causal or pleiotropic effects, rather than coincidental overlap of signal (as indicated by HEIDI heterogeneity test P>0.009) (**Table S7**). Secondly, we adopted a tissue-targeted eQTL approach by first identifying which of the 46 GTEx tissue types contain differentially-expressed genes that are enriched for AAM-associated variants. Five GTEx tissues, all in the central nervous system, were positively enriched for menarche-associated variants (Figure 2). Targeted assessment of these five enriched brain tissues, and also the hypothalamus which just missed significance for enrichment (P=9.8×10^−3^), using MetaXcan identified 205 genes whose expression was regulated by AAM-associated variants (**Table S8**). Of note, later AAM was associated with higher transcript levels of *LIN28B* (#147), *HSD17B12* (encoding Hydroxysteroid (17-Beta) Dehydrogenase 12; #250), and *NCOA6* (Nuclear receptor coactivator 6; #365).

**Figure 2.**
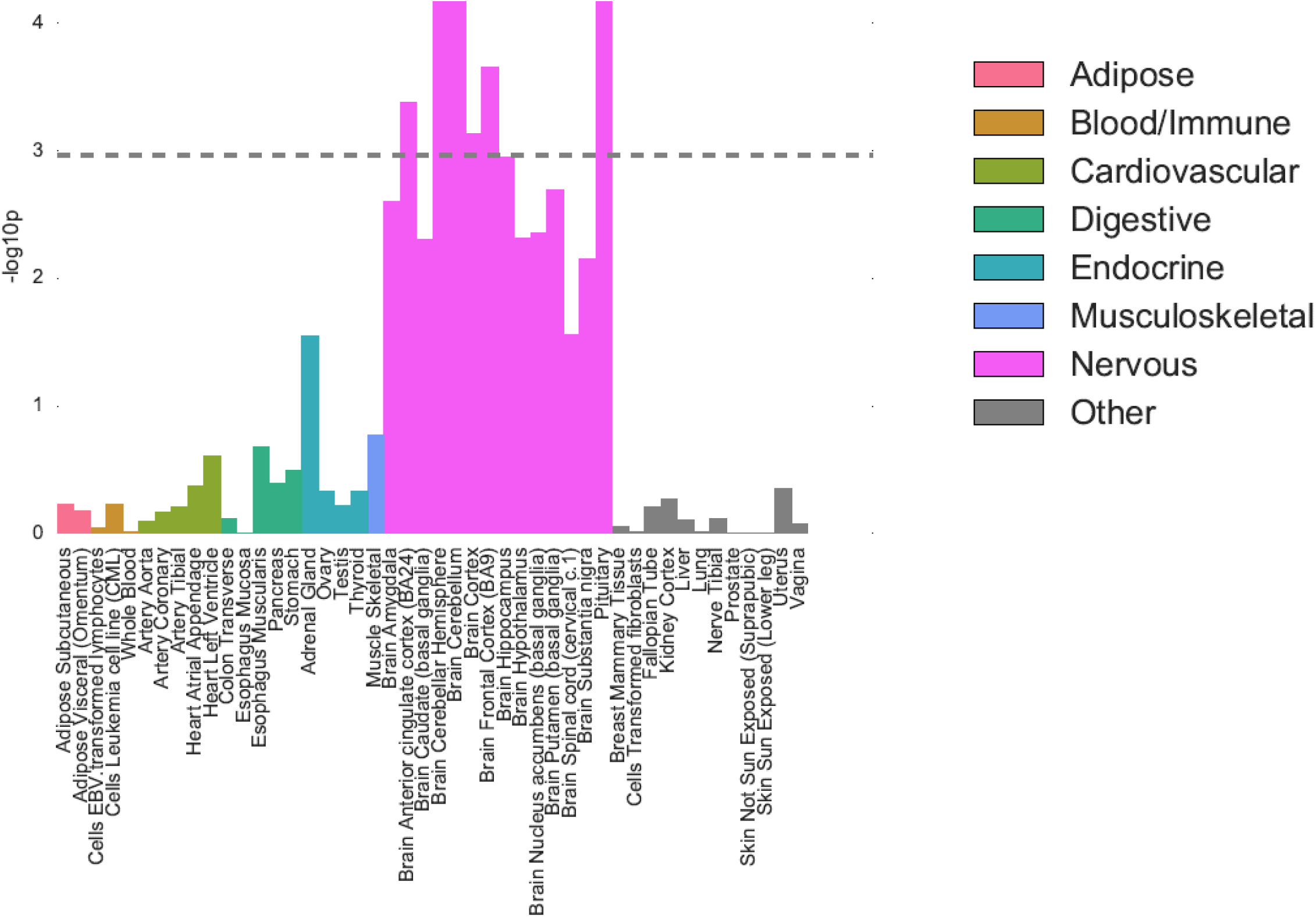
GTEx tissue enrichment using LD score regression

To identify possible distal causal genes, we interrogated reported Hi-C data to assess if any of the AAM loci are located in regions of chromatin looping^8^. 335 of the 389 loci were located within a topologically associating domain (TAD) – a defined boundary region containing chromatin contact points, each of which contained on average ~5 genes (**Table S9**). These included 22 of the 31 gene desert regions (nearest protein-coding gene >300kb), where TADs contained notable distal candidate genes such as *INHBA* (#158), *BDNF* (#248), *JARID2* (#128) and several gamma-aminobutyric acid receptors (#91). We also observed several regions where multiple independent menarche signals all reside within one TAD containing the same single gene – *RORB* (signal #200 intronic, signal #199 ~200kb downstream, #198 ~1.2Mb downstream), *THRB* (#67 intronic, #68 ~180kb upstream) and *TACR3* (#96 5’UTR, #97 ~25kb upstream, #98 ~133kb upstream and #95 ~263Kb downstream).

66 AAM signals were located in a specific contact point (between 5-25kb in size) within the 335 TADs, indicating a direct physical connection between these signals and a distal genomic region, on average ~320kb away. This included the previously reported example of the BMI-associated (and AAM-associated) *FTO* SNP and a distal *IRX3* promoter ~1Mb away (signal #326)^9^. The longest chromatin interaction observed was ~38.6Mb, where two distinct AAM signals located ~300kb apart (#206 and #207) were both in contact with the same distal genomic region ~38.6Mb away that contains only one gene: prostaglandin E synthase 2 (*PTGES2*).

## Transcription factor binding enrichment

To identify functional gene networks implicated in the regulation of AAM, we tested for enriched co-occurrence of menarche associations and predicted regulators within 226 enhancer modules combining DNaseI hypersensitive sites and chromatin states in 111 cell types and tissues. In total, we tested 2,382 transcription factor-enhancer module combinations. Sixteen transcription factor motifs were enriched for co-occurrence with AAM-associated variants within enhancer regions at study level significance (FDR<0.05) (**Table S10**). Furthermore, 5 of the 16 motif-associated transcription factors also mapped within 1Mb of an index AAM-associated SNP. These transcription factors included notable candidates; firstly, pituitary homeobox 1 (*PITX1*), is located within 50kb of genome-wide significant SNPs (~500kb from lead index #114). Secondly, *SMAD3*, a gene recently implicated in susceptibility to dizygous twinning^10^, is located within 600kb of an index SNP and its expression in several GTEx brain tissues is genetically correlated with AAM. Thirdly, *RXRB* is located within ~500kb of a novel index SNP (signal #133), and it represents the fifth (out of nine) retinoid-related receptor gene implicated by genome-wide significant AAM variants. This set now includes all three retinoid X receptor genes (*RXRA*, *RXRB* and *RXRG*), and retinoid-related receptor genes are the nearest gene to the index SNP at three AAM loci (*RXRA*, *RORA* and *RORB*).

## Pathway analyses

To identify other mechanisms that regulate pubertal timing, we tested all SNPs genome-wide for enrichment of AAM associations with pre-defined biological pathway genes. Ten pathways reached study-wise significance (FDR<0.05). Five pathways were related to transcription factor binding, and the other pathways were: peptide hormone binding, PI3-kinase binding, angiotensin stimulated signalling, neuron development and gamma-aminobutyric acid (GABA) type B receptor signalling (**Table S11**).

All of our previously reported custom pathways^3^ remained significant in this expanded dataset: nuclear hormone receptors (P=2.4×10^−3^); Mendelian pubertal disorder genes (P=1.9×10^−3^); and JmjC-domain-containing lysine-specific demethylases (P=1×10^−4^). Notably, new genome-wide significant signals mapped to lysine-specific demethylase genes: *JMJD1C* (signal #223), *PHF2* (#208), *KDM4B* (#347), *KDM6B* (#332), *JARID2* (#128), or to Mendelian pubertal disorder genes: *CYP19A1* (#307), *FGF8* (#230), *GNRH1* (#178) *KAL1* (#378), *KISS1* (#31), *NR5A1* (#215), and *NR0B1* (#379). The strongest menarche signal remains at *LIN28B*^3,11,12^, which encodes a key repressor of *let-7* miRNA biogenesis and cell pluripotency^13^. Transgenic *Lin28a/b* mice demonstrate both altered pubertal growth and glycaemic control^14^, suggesting that the *Lin28/let-7* axis could link puberty timing to type 2 diabetes susceptibility in humans. *let-7* miRNA targets are reportedly enriched for variants associated with type 2 diabetes^15^. We tested the same set of computationally-predicted and experimentally-derived mRNA/protein *let-7* miRNA targets^15^, and observed significant enrichment of AAM-associated variants at miRNA targets that are down-regulated by *let-7b* overexpression in primary human fibroblasts (**Table S16**, P_min_=1×10^−3^).

## Imprinted genes and parent-of-origin effects

We previously reported an excess of parent-of-origin specific associations for those AAM variants that map near imprinted genes, as defined primarily from animal studies^3^. Recent data from the GTEx consortium now allow a more systematic assessment of imprinted gene enrichment using genes defined from human transcriptome-wide analyses^16^. Consistent with our previous observations, imprinted genes were enriched for AAM-associated variants (MAGENTA P=4×10^−3^), with a concordant excess of parent-of-origin specific associations for the 389 index AAM variants (**Figure S2**, **Table S3**).

Systematic assessment of the 389 AAM gene regions in the Icelandic deCODE study revealed novel rare variants in two imprinted gene regions with robust parent-of-origin specific associations with AAM. Firstly, we identified a rare 5‘ UTR variant rs530324840 (MAF=0.80% in Iceland) in *MKRN3* that is associated with AAM under the paternal (P=6.4×10^−11^, β= −0.52 years) but not the maternal model (P=0.20, β=0.098) (Table 1 & **S12**). rs530324840 is by far the most significant variant at the *MKRN3* locus and is uncorrelated with our previously reported common variant rs12148769 at the same locus (r^2^ <0.001 in deCODE)^3^ (**Figure S3**). We note that the rare 5‘ UTR variant rs184950120 detected in the current GWAS meta-analysis also shows paternal-specific association in deCODE and, despite their near location (235bp from rs530324840), is uncorrelated to rs530324840 (r^2^<0.0001 in deCODE). No *MKRN3* loss-of-function variants were found in Iceland but five missense variants were observed and imputed. None of these five missense *MKRN3* variants detected in deCODE was associated with AAM in additive or parent-of-origin specific models (**Table S12**).

**Table 1:** Parent-of-origin specific associations between sequence variants at *MKRN3*, *DLK1 and MEG9* with age at menarche in Iceland (N=39,543).

| Marker | Position (hg38) | Allele | | Freq. A1 (%) | Region | Additive | | Maternal | | Paternal | | $P_{\text{mat vs. pat}}^2$ |
| --- | --- | --- | --- | --- | --- | --- | --- | --- | --- | --- | --- | --- |
| | | A1 | A2 | | | $P$ | $\beta^1$ | $P$ | $\beta^1$ | $P$ | $\beta^1$ | |
| rs530324840<sup>3</sup> | 15:23,565,461 | A | C | 0.80 | *MKRN3* | $4.4 \times 10^{-4}$ | -0.206 | 0.20 | 0.098 | $6.4 \times 10^{-11}$ | -0.523 | $1.3 \times 10^{-7}$ |
| rs184950120<sup>3</sup> | 15:23,565,696 | T | C | 0.26 | *MKRN3* | 0.010 | -0.265 | 0.98 | 0.003 | $1.5 \times 10^{-4}$ | -0.502 | 0.049 |
| rs12148769<sup>3</sup> | 15:23,906,947 | A | G | 10.1 | *MKRN3* | $5.8 \times 10^{-6}$ | -0.078 | 0.34 | -0.022 | $9.2 \times 10^{-8}$ | -0.120 | 0.0023 |
| rs138827001<sup>4</sup> | 14:100,771,634 | T | C | 0.36 | *DLK1* | $6.8 \times 10^{-6}$ | -0.387 | 0.88 | -0.018 | $4.7 \times 10^{-10}$ | -0.704 | $1.4 \times 10^{-4}$ |
| rs10144321<sup>4</sup> | 14:100,416,068 | G | A | 23.0 | *DLK1* | $5.6 \times 10^{-6}$ | -0.056 | 0.40 | -0.014 | $1.9 \times 10^{-7}$ | -0.084 | 0.0097 |
| rs7141210<sup>4</sup> | 14:100,716,133 | T | C | 38.2 | *DLK1* | 0.045 | 0.021 | 0.15 | -0.021 | $2.3 \times 10^{-5}$ | 0.059 | 0.0004 |
| rs61992671<sup>5</sup> | 14:101,065,517 | A | G | 48.5 | *MEG9* | 0.0047 | -0.029 | $6.0 \times 10^{-8}$ | -0.077 | 0.27 | 0.015 | $1.9 \times 10^{-5}$ |

1. $\beta$ indicates the effect of allele A1 in years per allele.
2. $P$ -value for heterogeneity between paternal and maternal allele associations.
3. rs530324840 is a novel variant identified by the parent-of-origin specific analysis. rs184950120 is the rare variant identified by the meta-analysis. rs12148769 is the previously reported intergenic common signal (Ref. 3).
4. rs138827001 is a novel variant identified by the parent-of-origin specific analysis. rs10144321 and rs7141210 are previously reported common variants (Ref. 3).
5. rs61992671 is a suggestive novel parent-of-origin specific association signal.

The second novel robust parent-of-origin specific signal is indicated by a rare intergenic variant at the *DLK1* locus (rs138827001; MAF=0.36% in Iceland) that associates with AAM under the paternal model (P=4.7×10^−10^, β= −0.70 years) but not the maternal model (P=0.88, β= −0.018 years) (Table 1, **Figure S4**). rs138827001 is uncorrelated with the two previously reported common variants rs10144321 and rs7141210 at the *DLK1* locus (r^2^ <0.01 in Iceland) that both also showed paternal allele-specific associations^3^. At this locus, we observed a further common variant rs61992671 (MAF=48.5% in Iceland) 4.4kb upstream of the Maternally Expressed 9 (*MEG9*) gene (~300kb from *DLK1*) that was associated with AAM under the maternal model (P=6.0×10^−8^, β= −0.077 years) but not the paternal model (P=0.27, β=0.015 years). rs61992671 was uncorrelated (r^2^<0.05) with the two common signals identified in the meta-analysis (rs10144321 and rs7141210) and replicated with a consistent magnitude of effect in the our GWAS meta-analysis (additive model, P=5.1×10^−6^).

## Disproportionate genetic effects on early or late puberty timing

Family-based studies in twins have suggested age-related differences in the impacts of genetic and environmental factors on AAM^17^. To test for asymmetry in the genetic effects on puberty timing, we defined two groups of women in the UK Biobank study based on approximated quintiles for AAM – “early” (8-11 years inclusive, N=14,922) and “late” (15-19, N=12,290). Each group was compared to the same median quintile AAM reference group (age 13, N=17,717). Estimated genome-wide heritability was higher for early AAM (h^2^_SNP_=28.8%; s.e 2.3%) than late AAM (h^2^_SNP_=21.5%; se 2.5%, P-difference=0.03). Accordingly, 217/377 (57.7%) autosomal index SNPs had larger effect estimates on early than late AAM (binomial P=0.004 vs. 50% expected), and the aggregated effect of the 377 SNPs also differed between strata (P=2.3×10^−4^) (Figure 3, **Table S13**). These differences remained when matching the early and late AAM strata for sample size and phenotype ranges (**Table S14**).

**Figure 3.**
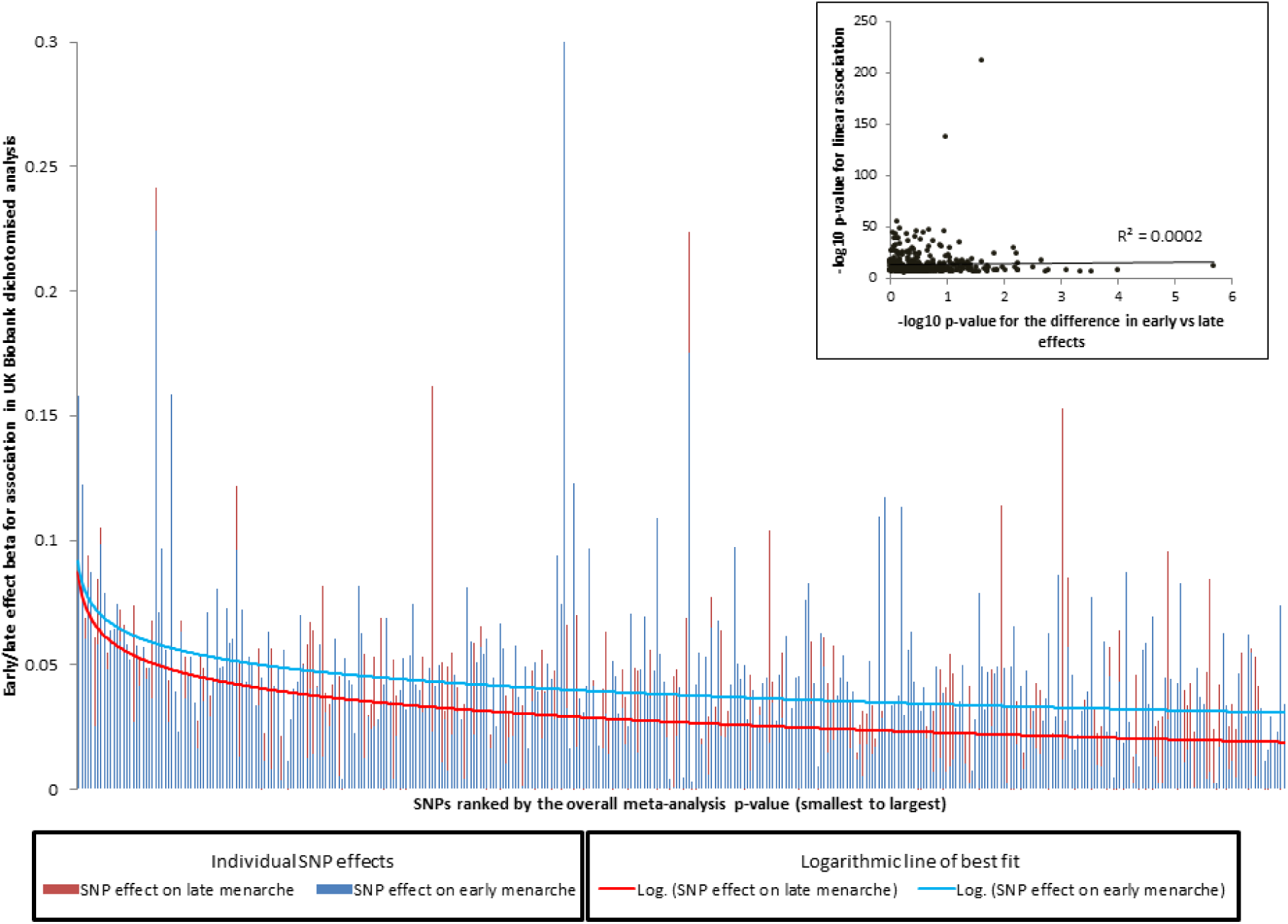
Stronger effects of age menarche-associated signals on early (blue) than late menarche (red) in women. The 377 index menarche-associated SNPs are ordered from smallest to largest p-value for their continuous associations with age at menarche. The Y-axis indicates the log-odds ratio for each SNP on early menarche (blue; ages 8–11 years inclusive) or late menarche (red; 15–19 years inclusive). The reference group are women with menarche at 13 years. **Insert** shows the –log_10_ p-values for the heterogeneity between the early and late menarche associations plotted against the –log_10_ p-value for the continuous age at menarche association.

**Figure 4.**
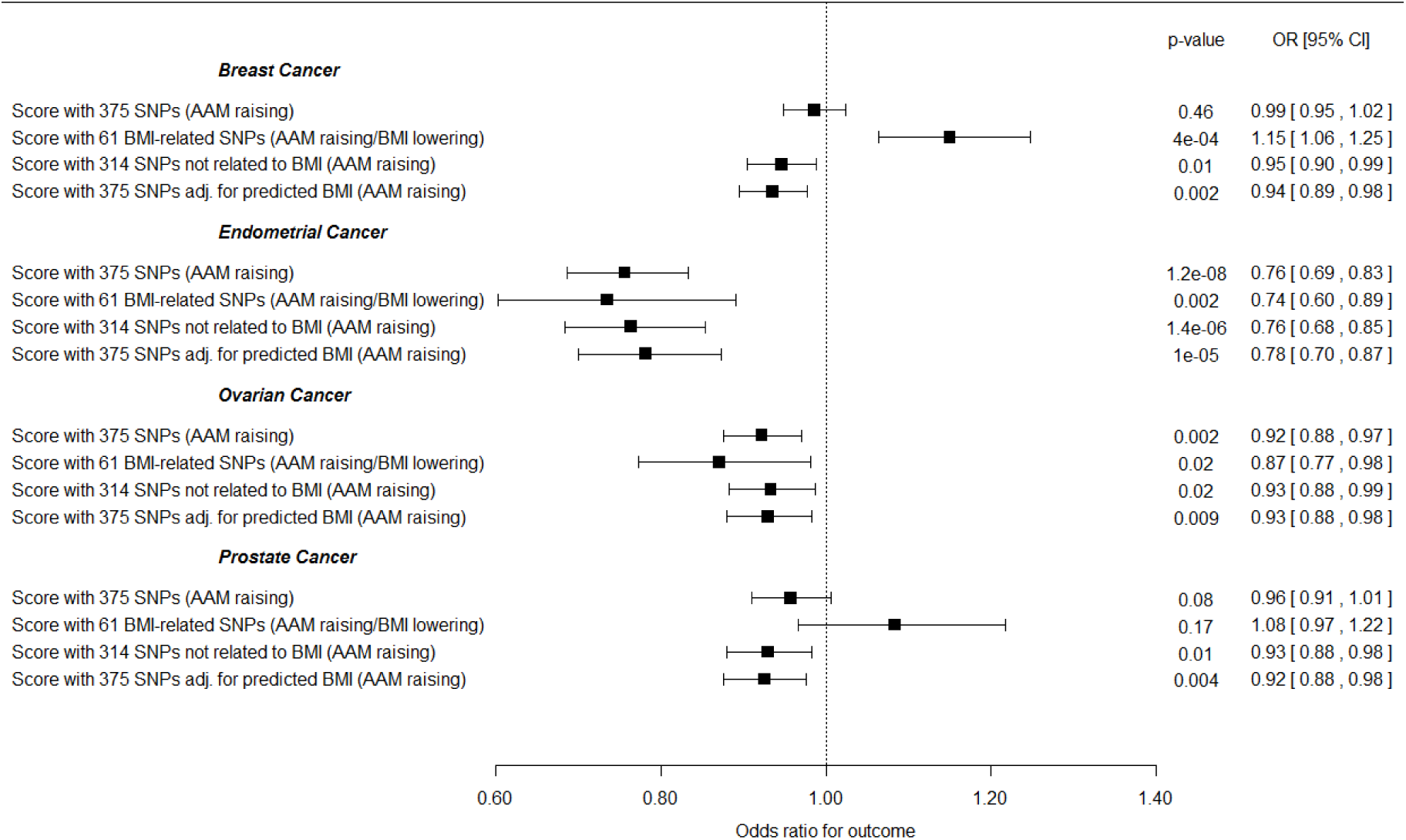
Effects of genetically-predicted age at menarche (AAM) on risks for various sex steroid-sensitive cancers, adjusted for the effects of the same AAM variants on BMI. AAM was predicted by all 375 autosomal AAM-associated SNPs, and models were adjusted for the genetic effects of the same AAM variants on BMI. Three further genetic score associations are shown as sensitivity analyses for each outcome: firstly, AAM predicted by the 314 AAM-associated SNPs that *were not* also associated with BMI in the BCAC iCOGs sample (at a nominal level of p<0.05); secondly, AAM predicted by the 61 AAM-associated SNPs that *were* also associated with BMI in this sample; finally, AAM predicted by all 375 autosomal AAM-associated SNPs (unadjusted for BMI).

In contrast, we observed the opposite pattern of disproportion in the genetic effects on male voice breaking in UK Biobank (“relatively early” N=2678, “relatively late” N=3566). Genome-wide heritability estimates tended to be higher for relatively late voice breaking (7.8%, s.e 1.2%) than for relatively early (6.9%, s.e 1.3%), and 227/377 (60.2%) index SNPs had larger effect estimates on relatively late than relatively early voice breaking (binomial P=4.3×10^−5^).

## BMI-independent effects of puberty timing on cancer risks

Traditional (non-genetic) epidemiological studies have reported complex associations between puberty timing, body mass index (BMI) and adult cancer risks. For example, large studies using historical growth records identified lower adolescent BMI and earlier puberty timing (estimated by the age at peak adolescent growth) as predictors of higher breast cancer risk in women^18,19^. Conversely, BMI is positively associated with breast cancer risk in postmenopausal women^20^. Furthermore, the strong inter-relationship between puberty timing and BMI limits the ability to consider their distinct influences on disease risks in traditional observational studies. Consistent with our previous report^5^, we observed a strong inverse genetic correlation between AAM and BMI (rg= −0.35, P=1.6×10^−72^). 39 AAM loci overlapped with reported loci for adult BMI^21^, yet even those AAM signals with weak individual associations with adult BMI still contributed to BMI when considered in aggregate: the 237 AAM variants without a nominal individual association with adult BMI (all P>0.05) were collectively associated with adult BMI (P=4.2×10^−9^) (**Figure S5**). This finding precludes an absolute distinction between BMI-related and BMI-unrelated AAM variants.

In Mendelian randomisation analyses, we therefore included adjustment for genetically-predicted BMI (as predicted by the 375 autosomal AAM variants) in order to assess the likely direct (i.e. BMI-independent) effects of AAM on the risks for various sex steroid-sensitive cancers (see **methods**). In these BMI-adjusted models, increasing AAM was associated with lower risk for breast cancer (OR=0.935 per year, 95% confidence interval: 0.894-0.977; P=2.6×10^−3^), and in particular with oestrogen receptor (ER)-positive but not ER-negative breast cancer (P-heterogeneity =0.02) (Figure 5, **Table S15**). Similarly, increasing AAM adjusted for genetically-predicted BMI was associated with lower risks for: ovarian cancer (OR=0.930, 0.880-0.982; P=9.3×10^−3^), in particular serous ovarian cancer (OR=0.917, 0.859-0.978; P=8.9×10^−3^); and endometrial cancer (OR=0.781, 0.699-0.872; P=9.97×10^−6^). Assuming an equivalent per-year effect of the current AAM variants on age at voice breaking, as we reported for the 106 previously identified AAM variants ^5^, we could also infer a protective effect of later puberty timing, independent of BMI, on lower risk for prostate cancer in men (OR=0.925, 0.876-0.976; P=4.4×10^−3^).

These findings were supported by sensitivity tests using sub-groups of AAM signals stratified by their individual associations with adult BMI. The ‘BMI-unrelated’ variant score (comprising 314 variants) supported a direct effect of AAM timing on breast cancer risk in women (OR=0.946, 0.904-0.988; P=1.3×10^−2^). In contrast, a score using only the 61 BMI-related AAM variants gave a significant result in the opposite direction (OR=1.15, 1.06-1.25; P=4.3×10^−4^) (**Table S15**), consistent with the recently reported inverse association between genetically-predicted BMI and breast cancer risk^22,23^. Further sensitivity tests (heterogeneity and MR-Egger tests) using the ‘BMI-unrelated’ variant score suggested that additional sub-pathways might link AAM to risk of ovarian cancer (MR-Egger Intercept P=0.036), but reassuringly these tests indicated no further pleiotropy (i.e. beyond the effects of BMI) in our analyses of breast, endometrial and prostate cancers (for all: I-square <23% and MR-Egger Intercept P>0.1) (**Table S15, Figure S6**).

## Discussion

In a substantially enlarged genomic analysis using densely imputed genomic data, we have identified 389 independent, genome-wide significant signals for AAM. In aggregate, these signals explain ~7.4% of the population variance in AAM, corresponding to ~25% of the estimated heritability. While assigning possible causal genes to associated loci is an ongoing challenge for GWAS findings, we adopted a number of recently described methods to implicate the underlying genes and tissues. 33 genes were implicated by non-synonymous variants and >200 genes were implicated by transcriptome-wide association in the five neural tissues enriched for AAM-associated gene activation. Transcriptome-wide association analyses also enabled the estimation of direction of gene expression in relation to AAM, notably indicating the likely delaying effect of *LIN28B* gene expression on AAM, which is consistent with inhibitory effects of this gene on developmental timing in animal and cell models^13,14^.

Our findings add to the growing evidence for a significant role of imprinted genes in the regulation of puberty timing^3^. In a recent family study, rare coding mutations (two frameshift, one stop-gained and one missense) in *MKRN3* were shown to cause central precocious puberty when paternally inherited^24^. None of those mutations were observed in the 8,453 whole-genome sequenced Icelanders in deCODE (median depth 32X). Taken together, three distinct types of variants at *MKRN3* appear to influence puberty timing when paternally inherited: (i) multiple rare loss-of-function mutations with large effects^24^ (ii) a common intergenic variant (rs530324840) with small effect, and (iii) two 5‘ UTR variants (rs184950120 and rs12148769) with intermediate allele frequencies (1 in 95 Icelandic women) and effects (~0.5 years per allele). Similarly, we found allelic heterogeneity at the imprinted *DLK1* locus where, as at *MKRN3*, a low frequency paternally-inherited allele conferred a substantial decrease in the age of puberty timing. At the same locus, maternal allele-specific association with an unrelated variant near to the maternally-expressed gene *MEG9* is consistent with multiple imprinting control centres at this imprinted gene cluster^25^.

The strong collective influence of the identified loci on AAM allowed informative stratification of AAM-associated variants in causal analyses to distinguish between BMI-related and BMI-unrelated pathways linking puberty timing to risk of sex steroid sensitive cancers. These findings were supported in BMI-adjusted models and, except for ovarian cancer, by additional tests for pleiotropy, and indicate causal influences of both lower adolescent BMI and earlier AAM on later cancer risks. The association between BMI and breast cancer risk is complex; directionally-opposing associations have been reported with adolescent and adult BMI, and with differing associations with pre- and post-menopausal breast cancer^18,19,20^. Recent Mendelian randomisation studies report a consistent protective effect of higher BMI on pre- and post-menopausal breast cancer^22,23^. Some studies have reported on the association between later puberty timing and lower risk of prostate cancer in men, but such data on puberty timing in men is scarcely recorded^26^. The influences of earlier puberty timing, independent of BMI, on higher risks of breast, ovarian and endometrial cancers in women, and prostate cancer in men, could be mediated by a longer duration of exposure to sex steroids. Alternatively, mechanisms that confer earlier puberty timing might also promote higher levels of hypothalamic-pituitary-gonadal axis activity, as exemplified by a variant in *FSHB* that confers earlier AAM, higher circulating follicle stimulating hormone concentrations in women, and higher susceptibility to dizygous twinning^10^.

We identified disproportionate effects of AAM variants on early or late puberty timing in a sex-discordant pattern. In females, variant effect estimates and heritability were higher for early versus late puberty timing, but the opposite was seen in males. These findings are concordant with clinical observations of sex-dependent penetrance of abnormal early and late puberty timing, even when accounting for presentation bias. Girls are more susceptible than boys to start puberty at abnormally young ages^27^, whereas boys are more susceptible than girls to have delayed onset of puberty^28^. These findings suggest some, yet to be unidentified, sex-specific gene-environment interactions. Future studies should systematically explore the potential influence of AAM-associated variants on rare disorders of puberty. In summary, our findings suggest unprecedented genetic complexity in the regulation of puberty timing and support new causal links with susceptibility to sex steroid-sensitive cancers in women and men.

## Online Methods

### GWAS meta-analysis for age at menarche in women

Each individual study tested SNPs using an additive linear regression model for association with age at menarche (AAM), including age at study visit and other study specific covariates. Insertion/deletion polymorphisms were coded as “I” and “D” for data storage efficiency and to allow harmonisation across all studies. Genetic variants and individuals were filtered on the basis of study specific quality control metrics. Association statistics for each SNP were then uploaded by study analysts for central processing. Study level results files were assessed following standardised quality control pipeline^29^, and results for each SNP were meta-analysed across studies using an inverse variance weighted model using METAL^30^ in a two stage process. Firstly, results from ReproGen consortium studies (**Table S1**) were combined and then filtered so that only those SNPs which appeared in over half of these studies were taken forward. Secondly, aggregated ReproGen consortium results were combined with data from the UK Biobank^31,32^ and 23andMe, Inc. studies^5^. Variants were only included in the final results file if they had results from at least two of these three sources, and a combined minor allele frequency (MAF) > 0.1%. We assessed potential inflation of test statistics due to sample relatedness and population stratification using LD score regression^33^. Here, an intercept value not significantly different from 1 indicates no such inflation, with a value over 1 indicating inflation.

A final list of index variants was first defined using a distance based metric, by which any SNPs passing the threshold of significance (P<5×10^−8^) within 1Mb of another significant SNP were considered to be located in the same locus. This list of signals was then further augmented using approximate conditional analysis in GCTA, using an LD reference panel from the UK Biobank study. Only secondary signals that were uncorrelated (r2<0.05) were included in the final list.

### Replication and parent-of-origin testing

Replication of identified hits was performed in an independent sample of 39,486 women of European ancestry from the deCODE study, Iceland. Main effects and parent-of-origin association testing was performed using the same methodology as previously reported^3,4^. The fraction of variance explained by a variant associating under the additive model was calculated using the formula 2 *f* (1−*f*) β_a_ ^2^, where *f* denotes the minor allele frequency of the variant and β_a_ is the additive effect. For variants associating under the recessive model, the formula *f*_*h*_ (1−*f*_*h*_) β_r_^2^ was used, where *f*_*h*_ denotes the homozygous frequency of the variant and β_r_ denotes the recessive effect. For variants associating under parent-of-origin models, fraction of variance explained was computed using the formulas *f* (1−*f*) β_m_^2^ for the maternal model and *f* (1−*f*) β_p_^2^ for the paternal model, where *f* denotes the minor allele frequency of the variant, β_m_ denotes the effect under the maternal model and β_p_ denotes the effect under the paternal model. Variance explained across multiple SNPs was calculated by summing the individual variances for all uncorrelated variants. We also estimate variance explained for top hits in UK Biobank using a combined allele score of all 377 autosomal genetic variants. Each individual variant was weighted using effect estimates derived from a meta-analysis excluding UK Biobank.

### Age at voice breaking in men

Data on male voice breaking were available from two sources. Firstly, the 23andMe, Inc. study recorded recalled age at voice breaking in a sample of 55,871 men, as previously described^5^. This was recorded as a quantitative trait into pre-defined 2-year age bins by online questionnaire in response to the question “How old were you when your voice began to crack/deepen?”^5^. Individual SNP effect estimates from the two year age bins were rescaled to 1 year estimates for both voice breaking and age at menarche as reported previously.

Age at voice breaking was also recalled in the UK Biobank study, as previously described^32^. This was recorded as a categorical trait: “younger than average”, “about average age”, “older than average”, “do not know” or “prefer not to answer” in response to the question “When did your voice break”. In separate models, the earlier or later voice breaking groups were compared to the average group (used as the reference group).

### Disproportionate effects on early or late puberty timing

Disproportionate effects on early or late puberty timing of AAM-associated SNPs were tested for AAM in UK Biobank. The distribution of AAM was divided into approximate quintiles, as previously reported^32^. Odds ratios for being in the youngest quintile (range 8-11) or the oldest (range 15-19) were compared to the middle quintile (age 13) as the reference, for each AAM-associated SNP and also for a combined weighted AAM-increasing allele score, with weights derived from a meta-analysis of all other studies except for UK Biobank. Sensitivity tests were performed by dividing UK Biobank individuals into broad strata based on birth year (before or after 1950) and geographic location (attendance at a study assessment centre in the North or South of the UK, as indicated by a line joining Mersey-Humber).

### Genetic correlation and genome-wide variance analysis

Genome-wide genetic correlations with adult BMI^21^ and voice breaking^5^ were estimated using LD score regression implemented in LDSC^33^. The total trait variance of all genotyped SNPs was calculated using Restricted Estimate Maximum Likelihood (REML) implemented in BOLT^34^. This was estimated using the same UK Biobank study sample in the discovery analysis, excluding any related individuals. The proportion of heritability explained by index SNPs was estimated by dividing the variance explained by the index SNPs, by the total variance explained by all genotyped SNPs genome-wide.

### Mendelian randomisation analyses

Individual genotype data on cancer outcomes were available from the Breast Cancer Association Consortium (BCAC) and Endometrial Cancer Association Consortium (ECAC). In addition, summary level results for ovary and prostate cancer were made available from the Ovarian Cancer Association Consortium (OCAC) and the Prostate Cancer Association Group to Investigate Cancer Associated Alterations in the Genome (PRACTICAL) consortium, respectively. Total analysed numbers were: 47,800 breast cancer cases and 40,302 controls, 4401 endometrial cancer cases and 28,758 controls, 18,175 ovarian cancer cases and 26,134 controls, and 20,219 prostate cancer cases and 20,440 controls (from the PRACTICAL iCOGS dataset).

We performed Mendelian randomisation analyses to assess the likely causal effects of puberty timing on the risks for various sex steroid-sensitive cancers. Hence, AAM was predicted by a weighted genetic risk score of all 375 autosomal AAM-associated SNPs, and genetically-predicted AAM was tested for association with each cancer in a logistic regression model. To avoid potential confounding by effects of the AAM genetic risk score on BMI, we performed BMI-adjusted analyses by including in models as a covariate the same AAM genetic risk score, but weighting each SNP for its effect on BMI (rather than on AAM) in the same study sample. Hence, we estimated the effect of genetically-predicted AAM controlling for genetically-predicted BMI by the same SNPs. As sensitivity tests, three further genetic score associations were performed for each cancer outcome: firstly, AAM predicted by the 314 AAM-associated SNPs that *were not* also individually associated with BMI in the BCAC iCOGs sample (at a nominal level of p<0.05); secondly, AAM predicted by the 61 AAM-associated SNPs that *were* also associated with BMI in this sample (i.e P<0.05); finally, AAM predicted by all 375 autosomal AAM-associated SNPs (unadjusted for BMI). To further consider pleiotropy, we tested for presence of heterogeneity between AAM-associated SNPs and analysed MR-Egger regression models ^35^.

### Pathway analyses

Meta-Analysis Gene-set Enrichment of variaNT Associations (MAGENTA) was used to explore pathway-based associations in the full GWAS dataset. MAGENTA implements a gene set enrichment analysis (GSEA) based approach, as previously described^36^. Briefly, each gene in the genome is mapped to a single index SNP with the lowest P-value within a 110 kb upstream, 40 kb downstream window. This P-value, representing a gene score, is then corrected for confounding factors such as gene size, SNP density and LD-related properties in a regression model. Genes within the HLA-region were excluded from analysis due to difficulties in accounting for gene density and LD patterns. Each mapped gene in the genome is then ranked by its adjusted gene score. At a given significance threshold (95th and 75th percentiles of all gene scores), the observed number of gene scores in a given pathway, with a ranked score above the specified threshold percentile, is calculated. This observed statistic is then compared to 1,000,000 randomly permuted pathways of identical size. This generates an empirical GSEA P-value for each pathway. Significance was determined when an individual pathway reached a false discovery rate (FDR) <0.05 in either analysis. In total, 3216 pathways from Gene Ontology, PANTHER, KEGG and Ingenuity were tested for enrichment of multiple modest associations with AAM. MAGENTA software was also used for enrichment testing of custom gene sets.

### Gene expression data integration

In order to identify which tissue types were most relevant to genes involved in pubertal development, we applied LD score regression^37^ to specifically expressed genes [method fully described in Finucane et al. in prep]. For each tissue, we ranked genes by a t-statistic for differential expression, using sex and age as covariates, and excluding all samples in related tissues. For example, we compared expression in hippocampus samples to expression in all non-brain samples. We then took the top 10% of genes by this ranking, formed a genome annotation including these genes (exons and introns) plus 100kb on either side, and used stratified LD score regression to estimate the contribution of this annotation to per-SNP AAM heritability, adjusting for all categories in the baseline model^37^. We computed significance using a block jackknife over SNPs, and corrected for 46 hypotheses tested at P=0.05.

To identify specific eQTL linked genes, we utilised two complementary approaches to systematically integrate publicly available gene expression data with our genome-wide dataset:

Summary Mendelian Randomization (SMR) uses summary-level gene expression data to map potentially functional genes to trait-associated SNPs^7^. We ran this approach against the publicly available whole-blood eQTL dataset published by Westra et al. ^6^, giving association statistics for 5,950 transcripts. A conservative significance threshold was set at P<8.4x10-6, in addition to a heterogeneity in dependent instruments (HEIDI) test statistic P>0.009 for any variants which surpass the main threshold.

MetaXcan, a meta-analysis extension of the PrediXcan method^38^, was used to infer the association between genetically predicted gene expression (GPGE) and age at menarche. PrediXcan is a novel gene-based data aggregation and integration method which incorporates information from gene-expression data and GWAS data to translate evidence of association with a phenotype from the SNP-level to the gene. Briefly, PrediXcan first imputes gene-expression at an individual level using prediction models trained on measured transcriptome datasets with genome-wide SNP data and then regresses the imputed transcriptome levels with phenotype of interest. MetaXcan extends its application to allow inference of the direction and magnitude of GPGE-phenotype associations with only summary GWAS statistics, which is advantageous when SNP-phenotype associations result from a meta-analysis setting and also when individual level data are not available. As input we utilized GWAS meta-analysis summary statistics for AAM, LD matrix from the 1000 Genomes project, and as weights, gene-expression regression coefficients for SNPs from models trained with transcriptome data (V6p) from the GTEx Project^39^. GTEx is a large-scale collaborative effort where DNA and RNA from multiple tissues were sequenced from almost 1,000 deceased individuals of European, African, and Asian ancestries. MetaXcan analyses were targeted to those tissue types with prior evidence of association with AAM (based on the GTEx enrichment analyses described above). The threshold for statistical significance was estimated using the Bonferroni method for multiple testing correction across all tested tissues (P<2.57x10^−6^).

### Motif enrichment testing and Hi-C integration

We identified transcription factors whose binding could be disrupted by AAM associated variants in enhancer regions by combining predicted enhancer regions across 111 human cell types and tissues with predicted motif instances of 651 transcription factor families as previously described^40^. We defined enhancer regions partitioned into 226 enhancer modules summarizing patterns of activity across 111 reference epigenomes by combining and clustering DNaseI hypersensitive sites and regions annotated with enhancer-like chromatin states^41^. We used the full GWAS summary statistics to identify a heuristic sub-genome-wide significance threshold, and used this threshold to identify modules enriched for AAM associated variants.

We used the enhancer modules to predict active regulators^42^, and then tested for enriched co-occurrence of predicted regulators and AAM associated variants surpassing our heuristic threshold within enhancer regions.

Hi-C interactions and topologically associated domains (TADs) defined from Rao et al. (multiple cell types combined, called at FDR < 10%) were intersected with identified SNPs based on genomic coordinates (build 37).

## Acknowledgements

This research has been conducted using the UK Biobank Resource. Full study-specific acknowledgements can be found in the online supplement.

